# The gut lactic acid bacteria metabolite, 10-oxo-*cis*-6,*trans*-11-octadecadienoic acid, suppresses inflammatory bowel disease in mice by modulating the NRF2 pathway and GPCR-signaling

**DOI:** 10.1101/2023.02.22.529495

**Authors:** Miki Ando, Kazuki Nagata, Ryuki Takeshita, Naoto Ito, Sakura Noguchi, Natsuki Minamikawa, Naoki Kodama, Asuka Yamamoto, Takuya Yashiro, Masakazu Hachisu, Gaku Ichihara, Shigenobu Kishino, Masayuki Yamamoto, Jun Ogawa, Chiharu Nishiyama

## Abstract

Various gut bacteria, including *Lactobacillus plantarum*, possess several enzymes that produce hydroxy fatty acids (FAs), oxo FAs, conjugated FAs, and partially saturated FAs from polyunsaturated FAs as secondary metabolites. Among these derivatives, we identified 10-oxo-*cis*-6,*trans*-11-octadecadienoic acid (γKetoC), a γ-linolenic acid (GLA)-derived enon FA, as the most effective immunomodulator, which inhibited the antigen-induced immunoactivation and LPS-induced production of inflammatory cytokines. The treatment with γKetoC significantly suppressed proliferation of CD4^+^ T cells, LPS-induced activation of bone marrow-derived dendritic cells (BMDCs), and LPS-induced IL-6 release from peritoneal cells, splenocytes, and CD11c^+^ cells isolated from the spleen. γKetoC also inhibited the release of inflammatory cytokines from BMDCs stimulated with poly-I:C, R-848, or CpG. Further in vitro experiments using an agonist of GPR40/120 suggested the involvement of these GPCRs in the effects of γKetoC on DCs. We also found that γKetoC stimulated the NRF2 pathway in DCs, and the suppressive effects of γKetoC and agonist of GPR40/120 on the release of IL-6 and IL-12 were reduced in *Nrf2*^*-/-*^ BMDCs. We evaluated the role of NRF2 in the anti-inflammatory effects of γKetoC in a dextran sodium sulfate-induced colitis model. The oral administration of γKetoC significantly reduced body weight loss, improved stool scores, and attenuated atrophy of the colon, in wild-type C57BL/6 and *Nrf2*^*+/-*^ mice with colitis. In contrast, the pathology of colitis was deteriorated in *Nrf2*^*-/-*^ mice even with the administration of γKetoC.

Collectively, the present results demonstrated the involvement of the NRF2 pathway and GPCRs in γKetoC-mediated anti-inflammatory responses.

## 1 Introduction

In the intestines, various secondary metabolites are produced by intestinal bacteria using food ingredient-derived materials as substrates. Several bacteria metabolites exert beneficial effects on the host body, such as short-chain fatty acids (FAs) produced from dietary fibers by *Clostridium*, which are involved in the maintenance of homeostasis and prevention of immune-related inflammatory diseases by modulating the function of both hematopoietic cells and non-hematopoietic cells. Although polyunsaturated FAs (PUFAs) are catalyzed by enzymes in host cells to achieve various bioactivities and their relationships with inflammatory diseases have been vigorously studied with a focus on the ω3/ω6 balance (1), a recent study revealed that PUFAs are also converted to derivatives, including hydroxy FAs, oxo FAs, conjugated FAs, and partially saturated FAs, through the catalysis of enzymes identified in the gut lactic acid bacterium, *Lactobacillus plantarum* (2). The PUFA metabolite 10-hydroxy-*cis*-12-octadecenoic acid (HYA), a hydroxy FA derived from linoleic acid (LA), regulates glucose homeostasis by activating GPR40 and GPR120, and increases resistance to obesity (3). The HYA-mediated activation of GPR40 has also been shown to accelerate the recovery of an impaired intestinal epithelial barrier (4) and disrupted gingival epithelial barrier (5). The metabolite 10-oxo-*cis*-12-octadecenoic acid (KetoA), an oxo FA derived from LA, enhances energy metabolism by activating TRPV1 in adipose tissue and exerts anti-obesity effects on the host body (6). KetoA is also involved in the regulation of host energy metabolism by accelerating adipocyte differentiation, adiponectin production, and glucose uptake through the activation of PPARγ (7). Another LA derivative 10-oxo-*trans*-11-octadecenoic acid (KetoC), an enon FA, was found to regulate the function of monocytes (8) and epithelial cells (9) via GPR120 signaling. Although accumulating evidence has demonstrated the beneficial effects of the bacteria metabolites of PUFAs on the host body, the roles of these metabolites in immune-related events remain unclear.

In the present study, we examined the effects of the bacteria-generated FAs on antigen (Ag)-induced immunoresponses and revealed that enon FAs suppressed the proliferation of T cells and the activation of dendritic cells (DCs). Detailed analyses focusing on 10-oxo-*cis*-6,*trans*-11-octadecadienoic acid (γKetoC), an enon FA derived from γ-linolenic acid (GLA), demonstrated that the release of inflammatory cytokines from DCs upon stimulation by TLR ligands was inhibited by γKetoC. To reveal the molecular mechanisms underlying the immunoregulatory effects of γKetoC, we investigated the involvement of GPCRs and the NF-E2-related factor 2 (NRF2) pathway. In addition, we utilized colitis model to wild-type (WT) mice and *Nrf2* knockout (KO) mice to evaluate the effects of γKetoC intake on the regulation of inflammatory responses *in vivo*.

## 2 Article types

Original Research.

## 3 Materials and Methods

### 3.1 Mice

C57BL/6 mice were purchased from Japan SLC (Hamamatsu, Japan). OT-II mice purchased from The Jackson Laboratory (USA) and previously generated *Nrf2*^*-/-*^ mice (10) were maintained on the C57BL/6 background. Mice were housed in a specific pathogen-free facility, and all animal experiments were performed in accordance with the guidelines of the Institutional Review Board of Tokyo University of Science. The present study was approved by the Animal Care and Use Committees of Tokyo University of Science: K22005, K21004, K20005, K19006, K18006, K17009, and K17012.

### 3.2 Cells

Bone marrow-derived DCs (BMDCs) generated as previously described (11), were stimulated with 100 ng/mL LPS (#L3024, Fujifilm Wako Chemicals Co., Ltd., Japan), poly-I:C (#P0913, Sigma-Aldrich), R-848 (#AG-CR1-3582-M005, AdipoGen), CpG (#tlrl-1826, InvivoGen). GW9508 (#10008907, Cayman Chemical, Ann Arbor, MI, USA) and YM-254890 (#257-00631, Fujifilm Wako Chemicals Co., Ltd.) were used as an agonist of GPR40 and GPR120 and an antagonist of Gq, respectively. Ovalbumin (OVA) peptide 323-339 (POV-3636-PI, Peptide Institute Inc., Osaka, Japan) was added to the culture medium of whole spleen cells prepared from OT-II mice to induce the antigen-presenting cell (APC)-dependent activation of CD4^+^ T cells. The MojoSort Mouse Naïve CD4^+^ T cell Isolation Kit (#480040, BioLegend), anti-CD3ε antibody (Ab) (clone 145-2C11, BioLegend), and anti-CD28 Ab (clone 37.51, BioLegend) were used for the isolation and stimulation of CD4^+^ T cells, respectively, as previously described (12). CD11c MicroBeads UltraPure, mouse (#130-125-835, Miltenyi Biotec) was used to isolate CD11c^+^ cells from the spleen.

### 3.3 Preparation of PUFA metabolites

Hydroxy, oxo, and enon FAs were prepared from LA, α-linolenic acid (ALA), and GLA, using the conversion enzymes isolated from *L. plantarum* AKU1009 (2). LA (#126-06571), and ALA (#122-05831) were purchased from Fujifilm Wako Chemicals and GLA (#L0152) from Tokyo Chemical Industry Co., Ltd. (Tokyo, Japan).

### 3.4 Enzyme-linked immunosorbent assay (ELISA)

The concentrations of mouse cytokines were measured using ELISA kits purchased from BioLegend (#431004 for IL-2, #431315 for IL-6, #430915 for TNF-α, and #431604 for IL-12p40, respectively).

### 3.5 Flow cytometry

CFSE (eBioscience Inc., San Diego, CA, USA) was used to monitor the proliferation of T cells. Surface MHC class II and CD86 on BMDCs were stained with anti-I-A/I-E-PerCP (clone M5/114.15.2, BioLegend) and anti-CD86-PE (clone GL-1, BioLegend), respectively. Fluorescence was detected by a MACS Quant Analyzer (Miltenyi Biotech) and analyzed with FlowJo (Tomy Digital Biology Co., Ltd., Tokyo, Japan).

### 3.6 Quantification of mRNA

The extraction of total RNA, synthesis of cDNA, and quantitative PCR were performed as previously described (13, 14). The nucleotide sequences of the primer sets used for qPCR are listed in Table I.

### 3.7 Western blot analysis

A Western blot analysis was performed with anti-NRF2 Ab (clone D1Z9C, Cell Signaling) and anti-β-actin Ab (clone AC-15, Sigma-Aldrich) as previously described (15).

### 3.8 Dextran sodium sulfate (DSS)-induced colitis

To induce colitis, mice were administered 2.5% (w/v) DSS (#160110, MP Biomedicals, Santa Ana, USA) in their drinking water. γKetoC (15 mg/kg/day) or vehicle (100 μl soybean oil) was orally administered using a sonde (#5202K, Fuchigami, Kyoto, Japan).

### 3.9 Statistical analysis

A two-tailed Student’s t-test was used for comparisons of two samples. To compare more than three samples, a one-way ANOVA-followed by the Tukey-Kramer multiple comparison test or Dunnett’s multiple comparison test was used. Area-under-curve (AUC) formatted data in DSS-induced colitis were calculated by GraphPad Prism 7.04. *P* values <0.05 were considered to be significant.

## 4 Results

### 4.1 Effects of bacteria metabolites of PUFAs on Ag-dependent responses in vitro

To examine the effects of bacteria metabolites of PUFAs on Ag-induced immunoresponses, we incubated OVA-stimulated OT-II spleen cells in the presence or absence of 50 μM of each metabolite for 48 h. The treatments with KetoC, αKetoC, γKetoA, and γKetoC markedly reduced the concentration of IL-2 in culture media, whereas those with HYA, αHYA, and γHYA did not (Fig. 1A). We then compared the suppressive effects of enon FAs on IL-2 production with those of the original PUFAs without conversion, and found that KetoC, αKetoC, and γKetoC significantly and dose-dependently suppressed IL-2 production, whereas apparent effects were not observed in LA, ALA, and GLA (Fig. 1B).

**Figure 1.**
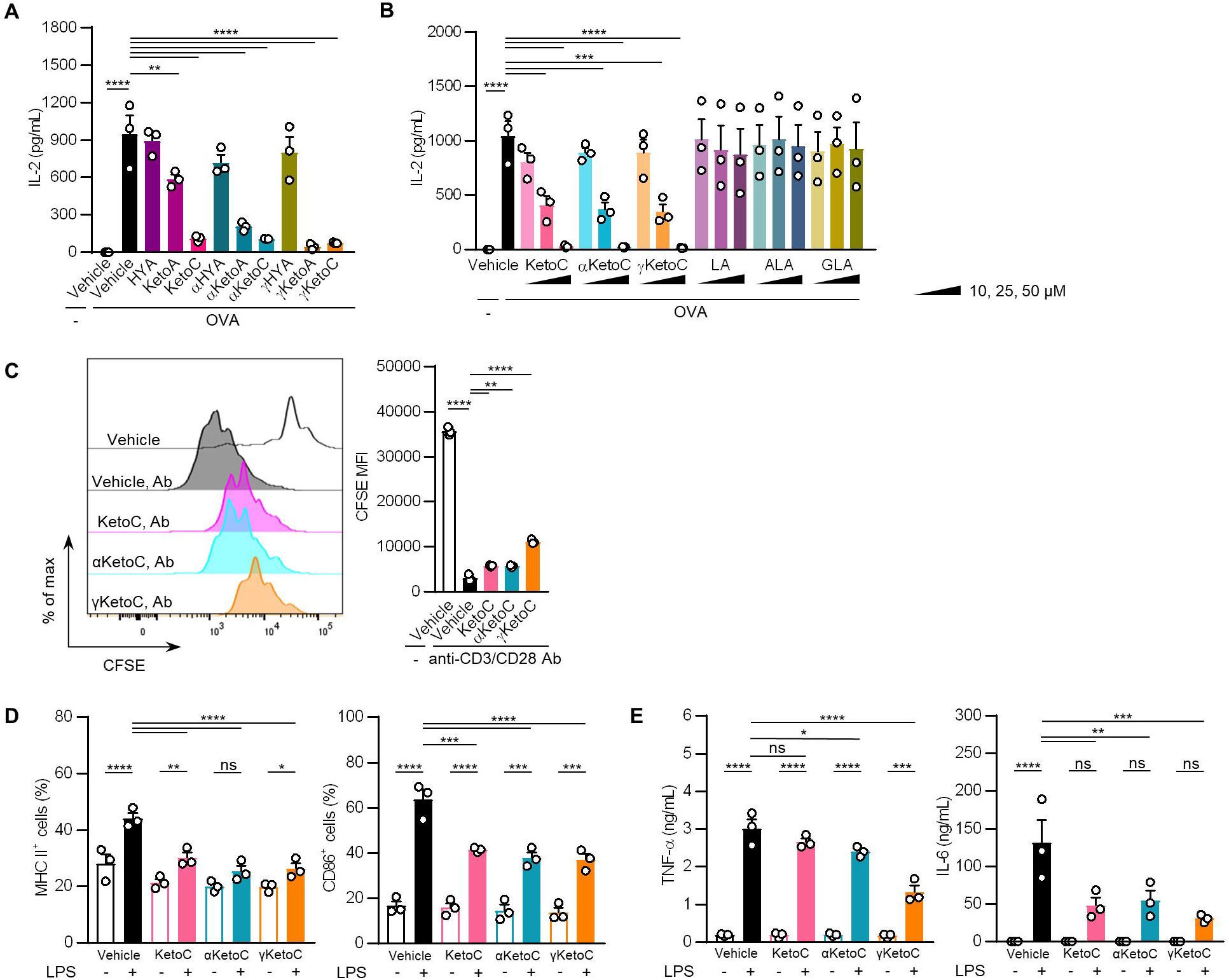
Effects of bacteria metabolites of PUFAs on the activation of T cells and DCs *in vitro*. **A**. and **B**. IL-2 concentrations in the culture media of splenocytes incubated in the presence or absence of OVA and FAs. In total, 1.0 × 10^5^/200 μL of OT-II spleen-derived single cell-suspended cells were stimulated by 2.5 μg/mL OVA with or without 50 μM bacteria metabolites of PUFAs or vehicle (ethanol) for 48 h (**A**). The indicated concentrations of enon FAs or their starting PUFAs were added to the culture media of OT-II spleen-derived cells with OVA during a 48-h incubation (**B**). **C**. The proliferation of CD4^+^ T cells stimulated with plate-coated anti-CD3 and anti-CD28 Abs. CD4^+^ T cells, which were isolated from the C57BL/6 spleen and were stained with CFSE, were incubated in Abs-coated dishes in the presence of 50 μM enon FAs for 72 h. **D**. Cell surface expression levels of MHC class II and CD86 in LPS-stimulated DCs. In total, 5.0 × 10^6^/2 mL of BMDCs were stimulated by 100 ng/mL LPS for 24 h in the presence or absence of 50 μM enon FAs. MFIs were shown as a ratio to that of LPS-stimulated BMDCs without FAs. **E**. Concentrations of cytokines in the culture media of LPS-stimulated DCs. In total, 5.0 × 10^6^/2 mL of BMDCs were stimulated by 100 ng/mL LPS for 24 h in the presence or absence of 50 μM enon FAs. Data represent the mean ± SEM of three independent experiments (**A, B, D, E**), and the mean ± SD of a typical data of triplicate samples from two independent experiments (**C**). The Dunnett’s test (**A**-**C**) and the Tukey-Kramer test (**D, E**) were used. \**p* < 0.05, \*\**p* < 0.01, \*\*\**p* < 0.005, \*\*\*\**p* < 0.0001, ns; not significant.

These results indicate that converted FAs carrying the enon structure acquired immunosuppressive effects, which were not observed in hydroxy FAs and were moderately induced in oxo FAs.

### 4.2 Suppressive effects of enon FAs on T cell proliferation and DC activation

To identify the cells in splenocytes that were regulated by the enon FAs, we examined the proliferation of T cells and the activation of DCs in the presence of enon FAs. The proliferation of naïve CD4^+^ T cells, which was induced by the treatment with plate-coated anti-CD3 and anti-CD28 Abs independent of APC, was suppressed by all three FAs at 50 μM (Fig. 1C). The pretreatment with 50 μM enon FAs also inhibited the up-regulation of MHC class II and CD86 on DCs (Fig. 1D) and the release of TNF-α and IL-6 from DCs (Fig. 1E) 24 h after the LPS stimulation.

These results demonstrate that enon FAs inhibited the activation of both of DCs and T cells, resulting in the suppression of Ag-induced IL-2 production in OT-II splenocytes.

### 4.3 γKetoC suppresses the wide spectrum of DC activation

To elucidate the mechanisms underlying the anti-inflammatory effects of enon FAs, we performed further analyses with a focus on γKetoC as the strongest suppressor among the three enon FAs. We confirmed that γKetoC significantly suppressed the LPS-induced production of IL-6 in peritoneal cells and whole leukocytes isolated from the spleen (Fig. 2A). The inhibition of IL-6 production by γKetoC was also observed in CD11c^+^ cells purified from the spleen (Fig. 2B). The effects of γKetoC on the activation of DCs indued by various stimulants were examined by using poly-I:C, R-848, and CpG. The measurement of cytokine concentrations revealed that the production of TNF-α, IL-6, IL-12p40 from BMDCs, which were induced by a stimulation via TLR3, TLR7/8, or TLR9, were significantly suppressed by γKetoC (Fig. 2C).

**Figure 2.**
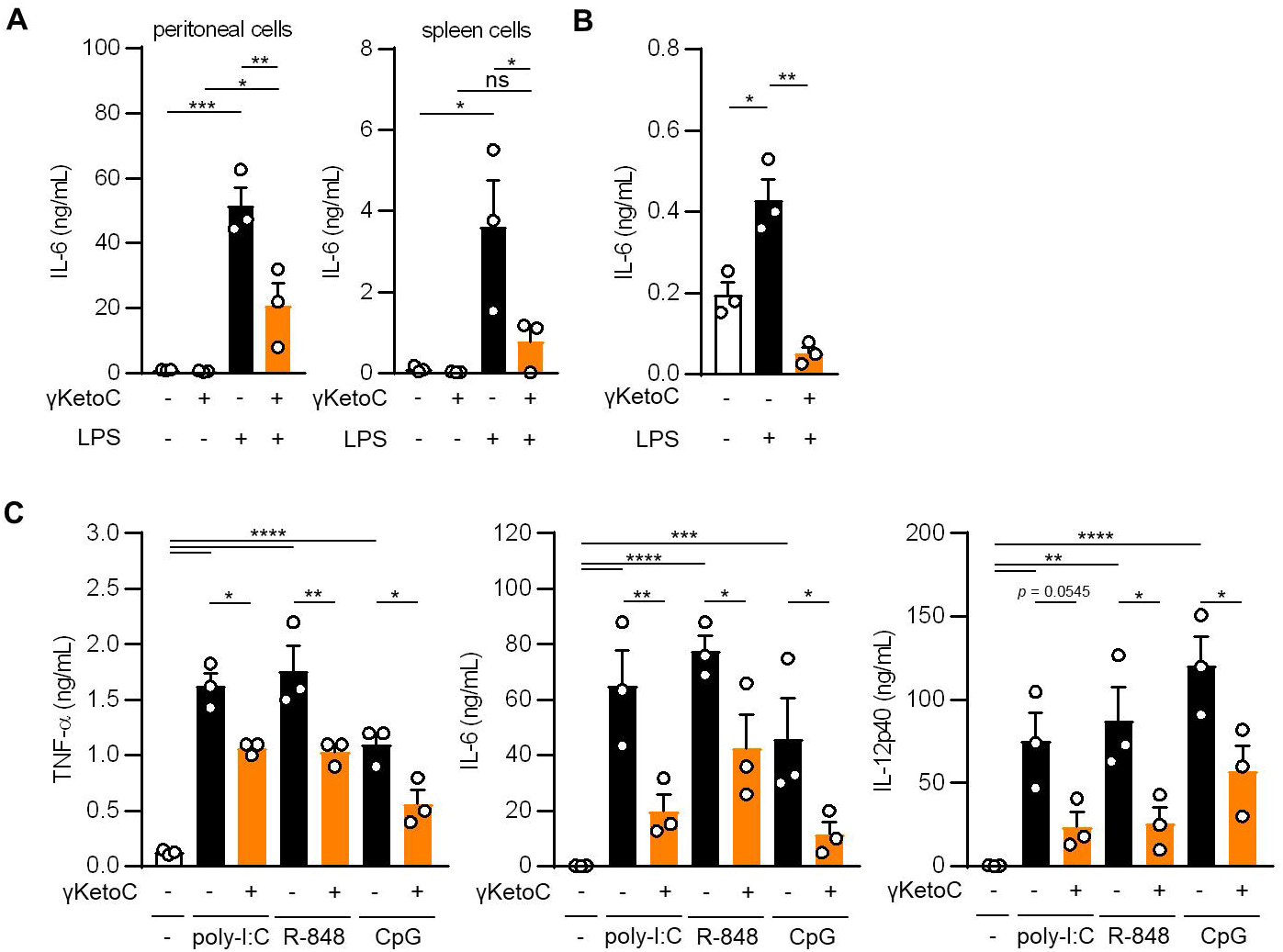
The suppressive effects of γKetoC on inflammatory cytokine producing cells stimulated with various TLR ligands. **A** and **B**. IL-6 release from LPS-stimulated peritoneal cells (**A** left), spleen cells (**A** right), and CD11c^+^ cells isolated from the spleen (**B**) was reduced by a treatment with γKetoC. Spleen cells (5.0 × 10^6^/mL), peritoneal cells (4.0 × 10^5^/mL), and CD11c^+^ cells (1.0 × 10^5^/200 μL) were stimulated with 100 ng/mL LPS for 24 h with or without 50 μM γKetoC. **C**. Concentrations of cytokines in the culture media of PAMPs-stimulated BMDCs. In total, 5.0 × 10^5^/500 μL of BMDCs were stimulated by 25 μg/mL poly-I:C, 1 μg/mL R-848, or 1 μg/mL CpG, for 24 h in the presence or absence of 50 μM enon FAs. Data represent the mean ± SEM of three independent experiments (**A, B, C**). The Tukey-Kramer (**A, C**) and Dunnett’s (**B**) multiple comparison test were used. \**p* < 0.05, \*\**p* < 0.01, \*\*\**p* < 0.005, \*\*\*\**p* < 0.001, ns; not significant.

### 4.4 Roles of Gq-GPCRs in the suppressive effects of γKetoC on the DC activation

Measurements of the mRNA levels of cytokines in LPS-stimulated BMDCs revealed that the inhibitory effects of γKetoC on transactivation was marked in the *Il12b*, and significant in *Il6* and *Tnf* (Fig. 3A). To clarify the molecular mechanisms by which γKetoC inhibited the PAMPs-induced transactivation of inflammatory cytokine genes in DCs, we first took notice of GPCRs based on the observation obtained in previous studies including ours. Briefly, KetoC inhibited the LPS-induced activation of the monocyte cell line RAW264.7 with binding to GPR120 (8), and an agonist of Gq-GPCR mimicked inhibitory effects of γKetoC on LPS-induced IL-6 production in BM macrophages (16). As in a previous report showing that GPR120 is expressed in adipocytes, macrophages, and DCs (17), GPR120 mRNA was detected in the BMDCs generated under our experimental conditions (data not shown). To examine the involvement of GPR120 in the γKetoC-mediated suppression of DCs, we treated BMDCs with GW9508, an agonist common to GPR40/GPR120, and revealed that GW9508 inhibited the LPS-induced release of inflammatory cytokines in a dose-dependent manner (Fig. 3B), suggesting that the stimulation of GPR120 suppressed the LPS-induced activation of DCs. Furthermore, the suppressive effects of γKetoC and GW9508 on LPS-induced production of TNF-α were abrogated by the pretreatment by YM-254890, an antagonist of Gq-GPCR, whereas YM-254890 did not alter the production of IL-6 and IL-12p40 (Fig. 3C).

**Figure 3.**
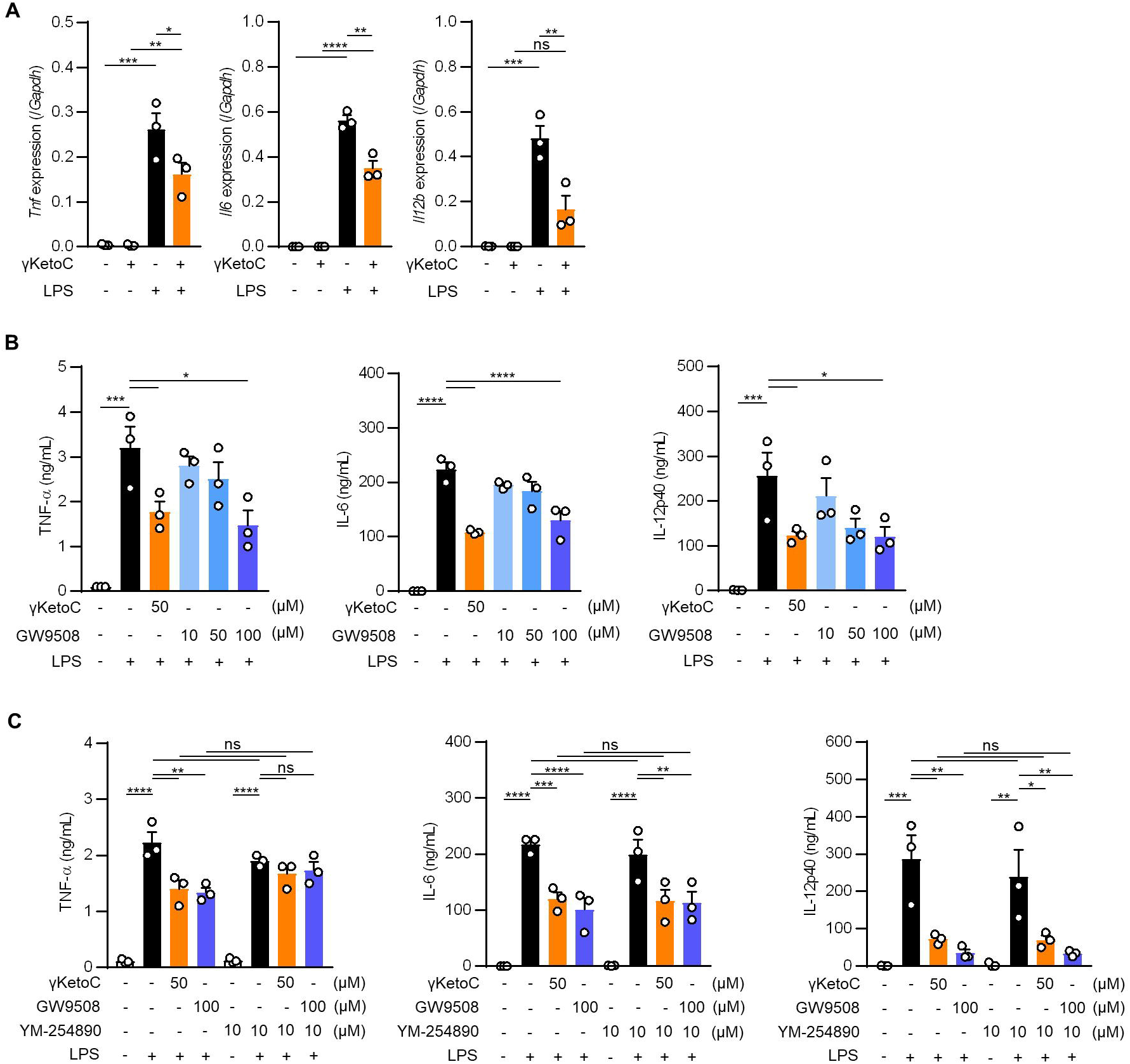
Roles of Gq-GPCR-signaling in the effects on DCs. **A**. mRNA levels of cytokine genes in LPS-stimulated DCs. 1.0 × 10^6^/mL of BMDCs were stimulated by 100 ng/mL LPS for 4 h in the presence or absence of 50 μM enon FAs. **B** and **C**. The amount of TNF-α, IL-6, and IL-12p40 released from LPS-stimulated DCs in the presence of the indicated concentrations of γKetoC or a Gq agonist (**B**) and those with a Gq antagonist (**C**). BMDCs pretreated in the presence or absence of the indicated concentrations of γKetoC, GW9508, and/or YM-254890 for 24 h, were cultured with or without LPS for an additional 24 h. Data represent the mean ± SEM of three independent experiments (**A, B, C**). The Tukey-Kramer test (**A, C**) and Dunnett’s test (**B**) was used. \**p* < 0.05, \*\**p* < 0.01, \*\*\**p* < 0.005, \*\*\*\**p* < 0.001, ns; not significant.

### 4.5 Involvement of the NRF2 pathway in the γKetoC-mediated suppression of DCs

Above-mentioned result indicating a partial involvement of GPCRs in the anti-inflammatory effects of γKetoC prompted us to analyze NRF2, a master transcription factor of antioxidant responses, as the other target of γKetoC. Previous studies reported that KetoC induced the expression of the antioxidant-related genes through the activation of NRF2, in the hepatic cell line HepG2 (18), and epithelial cell line Epi4 (9). In addition, a NRF2 deficiency enhanced the expression of IL-12p40 in stimulated DCs (14, 19). Therefore, to confirm whether γKetoC induced an antioxidant response via the activation of NRF2 in DCs, we examined NRF2 protein levels in γKetoC-treated DCs using Western blotting. The expression of NRF2 in BMDCs peaked at 1 h after the addition of γKetoC (Fig. 4A). The mRNA levels of *Hmox1*, a target gene of NRF2, were also increased in γKetoC-treated DCs (Fig. 4B). Furthermore, the suppressive effects of γKetoC on the LPS-induced IL-6 and IL-12p40 production were attenuated in *Nrf2*^*-/-*^ DCs (Fig. 4C). The effects of GW9508 on the production of IL-6 and IL-12p40 were also abrogated by the NRF2 deficiency (Fig. 4D). On the other hand, the deficiency of NRF2 did not affect the production levels of TNF-α (Fig. 4C, D).

**Figure 4.**
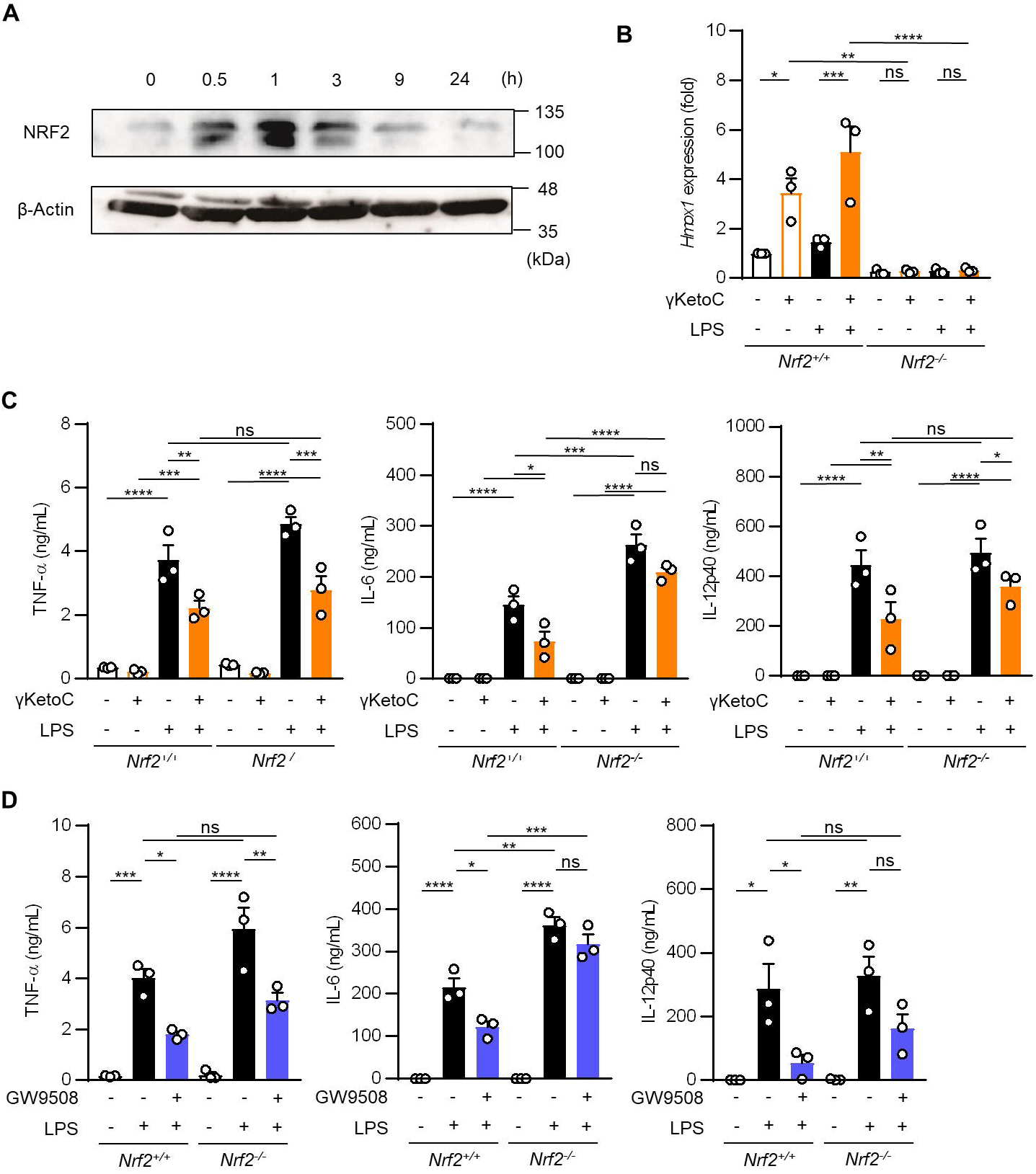
Involvement of the NRF2 pathway in the suppressive effects of γKetoC on DCs. **A**. NRF2 protein levels in γKetoC-treated DCs. BMDCs were cultured in the presence of 50 μM γKetoC for the indicated times, and aliquots of the whole cell lysate containing 10 μg of protein were applied to each lane of SDS-PAGE for Western blotting. **B**. mRNA levels of *Hmox1* in BMDCs derived from NRF2 deficient mice (*Nrf2*^-/-^) and its control (*Nrf2*^+/+^). 1.0 × 10^6^/mL of BMDCs were stimulated by 100 ng/mL LPS for 24 h in the presence or absence of 50 μM γKetoC. **C** and **D**. The amounts of cytokines released from NRF2-deficient DCs (*Nrf2*^-/-^) and their control DCs (*Nrf2*^+/+^). BMDCs derived from *Nrf2*^*+/+*^, or *Nrf2*^*-/-*^ mice, which were pretreated with or without 50 μM γKetoC (**C**) or 100 μM GW9508 (**D**) for 24 h, were cultured in the presence or absence of 100 ng/mL LPS for an additional 24 h. Data represent the mean ± SEM of three independent experiments performed in triplicate (**B, C, D**). The Tukey-Kramer test was used. \**p* < 0.05, \*\**p* < 0.01, \*\*\**p* < 0.005, \*\*\*\**p* < 0.001, ns; not significant.

These results indicate that γKetoC stimulated the NRF2 pathway via GPR120 resulting in the suppression of LPS-induced transactivation of the *Il6* and *Il12b* genes and subsequent cytokine production in DCs.

### 4.6 Oral administration of γKetoC ameliorates DSS-induced colitis

We utilized a DSS-induced colitis model to examine the protective effects of γKetoC on inflammatory responses *in vivo*. In the first colitis experiment, wild-type C57BL/6J mice were orally administered γKetoC (Fig. 5A). Although significant effect of γKetoC intake was not observed in the loss of body weight (Fig. 5B), increases in the disease activity index (DAI) score was alleviated by the intake of γKetoC (Fig. 5C). Fibrosis-mediated atrophy of the colon in mice with colitis was also significantly reduced in γKetoC-treated mice (Fig. 5D). In the next experiment, we investigated the roles of NRF2 in the γKetoC-mediated amelioration of colitis by using *Nrf2*^*-/-*^ mice (Fig. 5E). The administration of γKetoC was initiated 4 days earlier than that in the first experiment and the results obtained revealed that the loss of body weight (Fig. 5F) and increases in the DAI score (Fig. 5G) were significantly suppressed by γKetoC in *Nrf2*^*+/-*^ mice, but not in *Nrf2*^*-/-*^ mice. Atrophy of the colon in *Nrf2*^*+/-*^ mice was significantly restored by the intake of γKetoC, with the length of the colon being similar with and without the administration of γKetoC in *Nrf2*^*-/-*^ mice (Fig. 5H).

**Figure 5.**
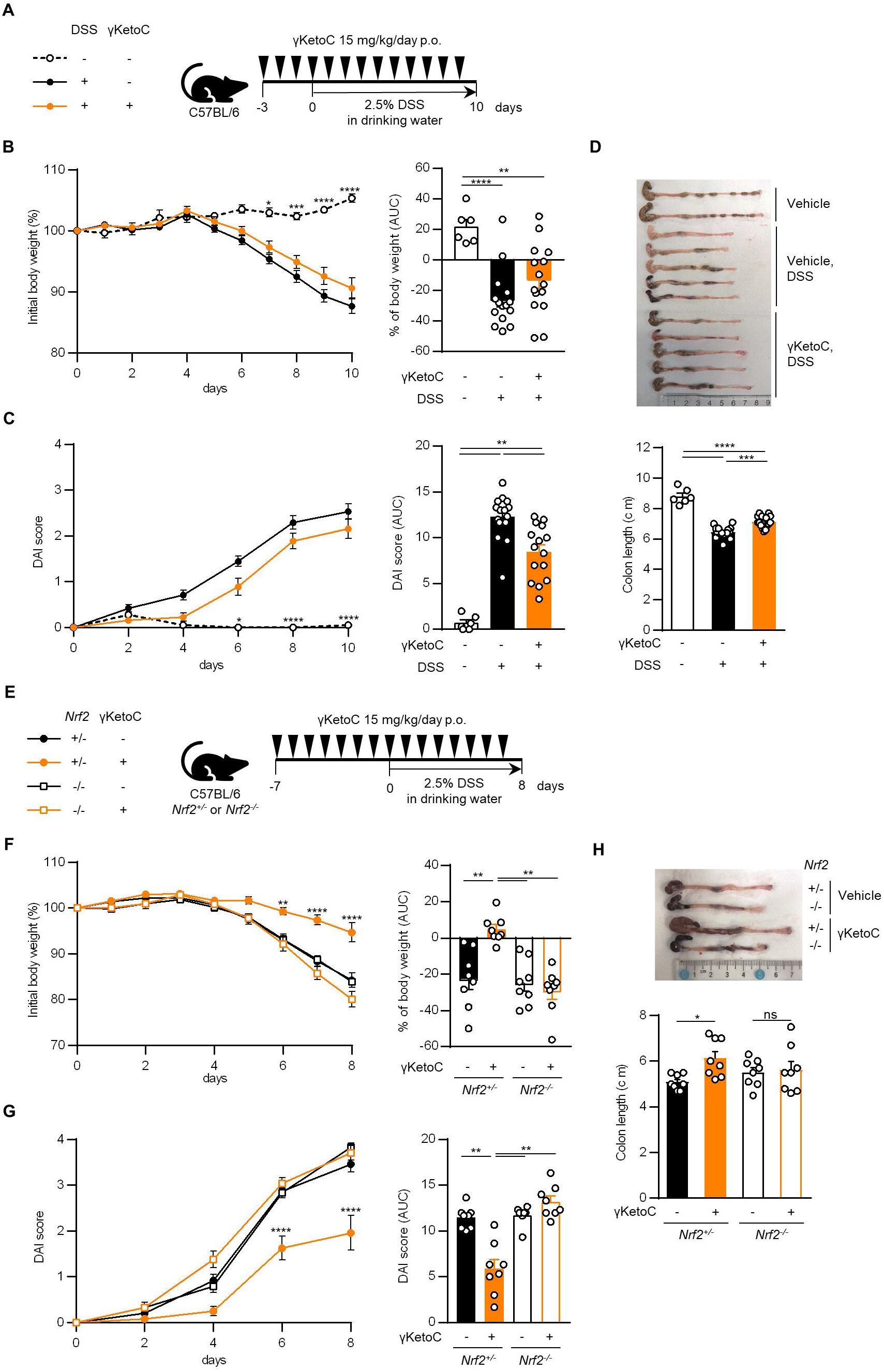
Effects of γKetoC on colitis in mice. **A**. Schematic of the oral administration schedule of γKetoC in the DSS-induced colitis model. C57BL/6J mice were orally administered 15 mg/kg/day of γKetoC in 100 μl soybean oil or vehicle. **B**. Percent body weight change from that measured on day 0 (left), and percent body weight change from day 0 to day 10 in area-under-curve (AUC) format (right). **C**. Disease activity index (DAI) scores (left), and DAI changes in AUC format (right). **D**. Images (top) and length (bottom) of the large intestine. DSS-; without the DSS treatment (n=6), DSS+; with the DSS treatment (n=15), DSS+γKetoC; administration of γKetoC with the DSS treatment (n=15) (**A** -**D**). **E**. Schematic of the schedule of administration of γKetoC (15 mg/kg/day) to colitis induced-*Nrf2* gene targeted mice. **F**. Percent body weight changes (left) and its AUC format (right). **G**. DAI score (left) and its AUC format (right). **H**. Atrophy levels of the colon. *Nrf2*^*+/-*^ DSS+; colitis-induced *Nrf2*^*+/-*^ mice (n=8), *Nrf2*^*+/-*^ DSS+γKetoC; γKetoC-treated colitis-induced *Nrf2*^*+/-*^ mice (n=8), *Nrf2*^*-/-*^ DSS+; colitis-induced *Nrf2*^*-/-*^ mice (n=8), *Nrf2*^*-/-*^ DSS+γKetoC; γKetoC-treated colitis-induced *Nrf2*^*-/-*^ mice (n=8) (**D** - **G**). Data are shown as the mean ± SEM. \**p* < 0.05, \*\**p* < 0.01, ns; not significant. The Tukey-Kramer test was used.

## 5 Discussion

The gut microbiota metabolizes food ingredients, and the resulting compounds exert beneficial effects on homeostasis in the host body. PUFAs, which are positively associated with inflammatory diseases depending on the amount consumed and the ω3/ω6 ratio, were recently shown to be modified by the enzymes of gut bacteria (2). Although previous studies demonstrated the useful effects of the bacteria metabolites of PUFAs on host health, particularly the attenuation of metabolic disorders (3, 6, 7), their effects on immunoresponses remain unclear.

The present results revealed that enon FAs suppressed Ag-mediated immunoresponses, which were not observed for their precursors, namely, LA, ALA, and GLA, or hydroxy FAs. Further analyses with a focus on γKetoC indicated that γKetoC suppressed the release of inflammatory cytokines from LPS (and other PAMPs)-stimulated DCs, whole splenocytes, and peritoneal cells. KetoC has been shown to inhibit the expression of inflammatory cytokines in LPS-stimulated RAW264.3 cells, and this was mitigated by a GPR120 antagonist, but not a GPR40 antagonist (8). In contrast to GPR40, which is a receptor for long-chain FAs as well as GPR120, but is highly expressed in the pancreas and liver and is involved in metabolism, GPR120 has an anti-inflammatory role as an receptor for ω3 FAs (17). Since HYA, which activates GPR40 (3-5), did not suppress the production of IL-2 by OVA-stimulated OT-II splenocytes in the present study (Fig. 1A), GPR40 might not play a prominent role in the regulation of inflammatory responses by immune-related cells. Based on the result showing that GW9508, a common agonist of GPR40 and GPR120, also reduced cytokine production by DCs, we speculate that GPR120 is involved in the anti-inflammatory effects of γKetoC as its receptor; however, we need to confirm this hypothesis in further experiments using a specific antagonist, siRNA, or knockout (KO) mice. γKetoC, KetoC, and αKetoC are categorized as ω7, ω7, and ω3, respectively. The structure of a FA required for ligand activity against GPR120 may not be the location of the unsaturated bond, but rather other factors, which were increased in enon FAs. If the enon structure is essential for binding to GPR120, metabolism by bacteria confers anti-inflammatory effects on dietary PUFAs.

In a SV40-T-transformed human gingival epithelial cell line, KetoC induced ERK phosphorylation and the subsequent activation of the NRF2 pathway via GPR120 (9). Under our experimental conditions, the GPR120 agonist did not induce *Hmox1* transactivation in DCs (data not shown), whereas the suppressive effects of the GPR120 agonist on cytokine production in DCs were reduced by a NRF2 deficiency. These results suggest that γKetoC activated both the NRF2 pathway and GPR120 in DCs and also that the NRF2 pathway might modulate GPR120 activity, whereas the stimulation of GPR120 did not induce an antioxidant response in DCs.

The present study suggested that the suppressive effects of γKetoC on TNF-α production were mediated by Gq-GPCR (probably GPR120), and the effects of γKetoC on production of IL-6 and IL-12p40 was largely dependent on NRF2. These differences might reflect the promoter specific roles of NRF2 and/or GPR120. A vigorous study revealed the direct binding of NRF2 to the *Il6* promoter inhibits the transcription of the *Il6* gene in macrophages (20). It is also known the anti-inflammatory effects of DHA is caused by the GPR120-dependent activation of β-arrestin (17). Further detailed analyses regarding GPR120-mediated signal transduction and NRF2 recruitment toward chromosomal DNA in γKetoC-treated DCs are required to uncover the molecular mechanisms of anti-inflammatory effects of γKetoC. In addition, the roles of γKetoC in T cell function were still unclear in the present study, even though DC-independent proliferation of T cells were significantly suppressed by γKetoC. We need to perform the experiments investigating the effects of γKetoC on various immuno-related cells, to clarify overall immunoregulatory effects of γKetoC.

γKetoC increased NRF2 protein and *Hmox1* mRNA levels in DCs, and NRF2 deficiency reduced the anti-inflammatory effects of γKetoC both *in vitro* and *in vivo*. NRF2 is a ubiquitous transcription factor, and *Nrf2* KO mice exhibit severe inflammation in various immune-related diseases, including contact hypersensitivity, autoimmune disease, colitis, and psoriasis (21-26). Therefore, γKetoC and other enon FAs have the potential to prevent and/or treat immune-related diseases. In addition, since we recently demonstrated that γKetoC suppressed osteoclast development and macrophage activation (16), it may also attenuate rheumatoid arthritis.

The present study showed that several bacteria metabolites of PUFAs, particularly enon FAs, were involved in the regulation of immunoresponses, which were not observed for their precursors. The NRF2 pathway and GPR120, both of which play important roles in anti-inflammatory responses, appear to be involved in the effects of γKetoC. The intake of γKetoC ameliorated colitis in mice in a NRF2-dependent manner. Based on these results, we conclude that gut bacteria and their metabolites of PUFAs exert beneficial effects on immune homeostasis in the host body.

## 6 Data availability statement

The original contributions presented in this study are included in the article/supplementary material, further inquiries can be directed to the corresponding author.

## 7 Conflict of Interest

The authors declare that the research was conducted in the absence of any commercial or financial relationships that could be construed as a potential conflict of interest.

## 8 Author Contributions

Contribution: M.A., K.N., and R.T. performed experiments, analyzed data, and prepared figures; N.I., S.N., N.M., N.K., A.Y., and T.Y. performed experiments; M.H. analyzed data; G.I., S.K., M.Y., and J.O. provided essential experimental tools and materials; C.N. designed research and wrote the paper.

## 9 Funding

This work was supported by a Grant-in-Aid for Scientific Research (B) 23H02167 (CN) and 20H02939 (CN); a Research Fellowship for Young Scientists DC2 and a Grant-in-Aid for JSPS Fellows 21J12113 (KN); a Scholarship for a Doctoral Student in Immunology (from JSI to NI); a Tokyo University of Science Grant for President’s Research Promotion (CN); the Tojuro Iijima Foundation for Food Science and Technology (CN); a Research Grant from the Mishima Kaiun Memorial Foundation (CN); and a Research Grant from the Takeda Science Foundation (CN).

## 10 Acknowledgments

We thank the members of the Laboratory of Molecular Biology and Immunology, Department of Biological Science and Technology, Tokyo University of Science for constructive discussions and technical support. We greatly appreciate the consideration and support from Dr. Kimihiko Yasuda, Dr. Masako Yasuda, and the late Ms. Yayoi Yasuda.

## 12 Supplementary Material

None.

## Notes

### Competing Interest Statement

The authors have declared no competing interest.

### Summary of Updates

Ryuki Takeshita and Natsuki Minamikawa are added in the author list. Several data are added in Fig.s 2, 3, and 4.

